# The unique and enigmatic spirochete symbiont of latrunculid sponges

**DOI:** 10.1101/2024.05.23.595633

**Authors:** Samantha C. Waterworth, Gabriella M. Solomons, Jarmo-Charles J. Kalinski, Luthando S. Madonsela, Shirley Parker-Nance, Rosemary A. Dorrington

**Author notes:** **Address correspondence to:** Rosemary A. Dorrington.

## Abstract

Bacterial symbionts are critical members of many marine sponge holobionts. Some sponge-associated bacterial lineages, such as Poribacteria, SAUL, and Tethybacterales appear to have broad host ranges and associate with a diversity of sponge species, while others are more species-specific, having adapted to the niche environment of their host. Host-associated spirochete symbionts that are numerically dominant have been documented in several invertebrates including termites, starfish, and corals. However, dominant spirochete populations are rare in marine sponges, thus far only observed in *Clathrina clathrus* and various species within the Latrunculiidae family, where they are co-dominant alongside Tethybacterales symbionts. This study aimed to characterize these spirochetes and their potential role in the host sponge. Analysis of metagenome-assembled genomes from eight latrunculid sponges revealed that these unusual spirochetes are relatively recent symbionts and are phylogenetically distinct from other sponge-associated spirochetes. Functional comparative analysis suggests that the host sponge may have selected for these spirochetes due to their ability to produce terpenoids and/or possible structural contributions.

**IMPORTANCE:** South African latrunculid sponges are host to co-dominant Tethybacterales and Spirochete symbionts. While the Tethybacterales are broad-host range symbionts, the spirochetes have not been reported as abundant in any other marine sponge except *Clathrina clathrus*. However, spirochetes are regularly the most dominant populations in marine corals and terrestrial invertebrates where they are predicted to serve as beneficial symbionts. Here, we interrogated eight metagenome-assembled genomes of the latrunculid-associated spirochetes and found that these symbionts are phylogenetically distinct from all invertebrate-associated spirochetes. The symbiosis between the spirochetes and their sponge host appears to have been established relatively recently.

## INTRODUCTION

The development of symbiotic relationships with prokaryotes likely predates the emergence of marine sponges (phylum Porifera) during the Cambrian explosion ∼540 million years ago (1, 2) and these associations have played a critical role in the evolution of modern sponge taxa (3, 4). Bacterial symbionts have co-evolved with their host to perform specific, specialized services that promote the health and fitness of the host (5). The symbionts are involved in nitrogen, sulfur, and phosphorus cycling (6–9), carbon cycling, detoxification (10, 11) and in some cases, the production of bioactive secondary metabolites as chemical defenses against pathogens, predators, and competitors (12, 13). In return, the host provides its symbionts with a safe and nutrient-rich environment that promotes the fitness and survival of the symbiont (14). The taxonomic and functional diversity of sponge-associated microbiomes is generally host-specific, distinct from the surrounding water column, and acquired by recruitment and enrichment from the environment (5, 15, 16). However, there are a small number of specialized symbionts acquired by vertical inheritance from the parent sponge that are broadly distributed across phylogenetically distant sponge hosts (17, 18), including the Poribacteria, the “sponge-associated unclassified lineage” (SAUL), and the recently-discovered Tethybacterales symbionts (15, 19, 20).

The Tethybacterales represent a clade of cosmopolitan sponge-associated symbionts, comprising three families, namely the *Candidatus* Persebacteraceae, *Candidatus* Tethybacteraceae, and *Candidatus* Polydorabacteraceae (17, 20). As with the Poribacteria and Desulfobacteria, the Tethybacterales symbionts are present in phylogenetically diverse taxa that are primarily low-microbial abundance (LMA) sponge species but these bacteria have also been detected in some high-microbial abundance (HMA) species (17, 20). Characterization of metagenome-assembled genomes (MAGs) of different species of the three Tethybacterales families and their associated hosts also indicates that there were multiple acquisition events and that host adaptation and co-evolution began after each acquisition event (17).

Sponges of the family Latrunculiidae (Demospongiae, Poecilosclerida) are known to be prolific producers of cytotoxic pyrroloiminoquinone alkaloid compounds (21–26) with pharmaceutical potential (Reviewed in Kalinksi et al., 2022 (27)). It has recently been discovered that there are two chemotypes present in the *Tsitsikamma favus* and *Tsitsikamma michaeli* latrunculid sponges (21, 28). Latrunculids are LMA sponges with highly conserved microbiomes that are dominated by Tethybacterales and Spirochete taxa (22, 29). The *Tsitsikamma favus* microbiome is dominated by two sponge-specific bacterial species defined by their 16S rRNA gene sequence, clones Sp02-1 and Sp02-3. The Sp02-1 symbiont has been recently characterized (17) and is classified as *Ca.* Ukwabelana africanus, a member of the *Ca.* Persebacteraceae family within the Tethybacterales (17). The *Ca.* U. africanus symbiont is phylogenetically related to symbionts in sponges across multiple orders within the Demospongiae and may be involved in the reduction of nitrogen and sulfur in the sponge holobiont (17).

Unlike *Ca.* U. africanus (Sp02-1), the co-dominant spirochete (Sp02-3) is not representative of a globally distributed, broad-host range sponge symbiont. Spirochetes have been reported as minor members of several sponge microbiomes (30–32), but numerically dominant populations of spirochetes in sponges have only been reported in Latrunculiidae species endemic to the southeastern coast of South Africa, and the distantly related *Clathrina clathrus* (Calcarea, Clathrinida) collected by Neulinger and colleagues from the Adriatic Sea off the coast of Croatia (33). In addition, spirochetes, presumed to be symbionts, have been detected in the embryonic and larval cells of the marine sponge *Mycale laevis*, but their role is currently unknown (34, 35). Numerically dominant spirochete species are, however, present in several other marine invertebrates including sea anemones (36) and sea stars (37, 38) where decreased abundance of certain spirochete populations correlates with an increased incidence of disease (38). Spirochaeta symbionts are also commonly present as dominant populations in corals (39–42) and in termite guts (43), where they may be involved in the fixation of carbon or nitrogen (41). A recent study investigating the association between coral hosts and their associated microbiota found that Spirochaeta were most abundant in the coral skeleton, hypothesizing that they may be key members in coral skeletal environment due to their ability to fix carbon and nitrogen (44).

The aim of the present study was to understand the relationship between latrunculid sponges and the Sp02-3 spirochete symbiont. Here we report the characterization of eight spirochete MAGs from four *Tsitsikamma* sponge species and use comparative genomics to shed light on factors that may drive their conservation. Comparative analysis relative to publicly available genomes and MAGs of the Spirochaetaceae family suggests that the Sp02-3 spirochetes are distinct from all other sponge-associated spirochetes.

## RESULTS AND DISCUSSION

Previous studies identified two closely related spirochete species, Sp02-3 and Sp02-15, in the *T. favus* microbiome (22). Subsequently, the Sp02-3 symbiont was shown to be present in the microbiomes of other *Tsitsikamma* species and *Cyclacanthia bellae* (29). Our aim in this study was to characterize the genome of the Sp02-3 symbiont to better understand its role in the sponge holobiont.

### Survey of microbial communities in latrunculid sponges and other sponge species endemic to the South African coast

To survey the prevalence of spirochetes in sponge collected off the South African coastline, we clustered 16S rRNA gene fragment amplicons sourced from 155 marine sponges and 8 seawater samples into operational taxonomic units (OTUs) at a distance of 0.03 in mothur (45). These sponges were collected primarily from reefs within Algoa Bay, South Africa but also included samples from the Tsitsikamma National Park, the Amathole Marine Protected Area in the Indian Ocean, and the remote Bouvet Island in the Southern (Antarctic) Ocean (Table S1).

A total of 9711 OTUs were recovered from the 163 amplicon libraries. We identified spirochete OTUs with classifications from alignment of the OTUs against the SILVA and nr databases (Table S2). A total of 142 OTUs were classified within the Spirochaetota phylum, of which only 10 had an average abundance greater then 0.01% across all sponge specimens (Fig. 1A). OTU3 and OTU59 were most abundant in the *Tsitsikamma* and *Cyclacanthia* sponges. These OTUs were most closely related to spirochete 16S rRNA gene clones Sp02-3 and Sp02-15, previously identified in *T. favus* sponges (22). These two OTUs were present at low abundance in the *Latrunculia algoaensis* and *Latrunculia apicalis* sponge specimens (collected in Algoa Bay and the Southern Antarctic Ocean), as well as in some *Mycale* specimens and a single sympatric *Phorbus sp.* sponge (Fig. 1B).. As the *Mycale* specimens were found as encrusting species on the *Tsitsikamma favus* sponges, we cannot discount the possibility of contamination between these two species. As we have only a single *Phorbus* sp. representative, additional specimens will be required to determine the significance of these spirochete OTUs in this genus or whether this was a result of contamination during collection. These two OTUs were otherwise absent in all other non-latrunculid sponges collected from sympatric regions. The presence, albeit low, of OTU3 and OTU59 in the *L. apicalis* sponges collected just off of Bouvet Island (∼ 3000 km/ 1800 miles from Algoa Bay), and the presence of phylogenetically distinct spirochetes in sympatric non-latrunculid sponges of Algoa Bay would suggest that these Sp02-3 and Sp02-15 spirochetes are specifically associated with latrunculid sponges Spirochete OTUs OTU105 and OTU128 were relatively abundant in other sponges collected from the South African coast, and absent in latrunculid sponges, appeared more sporadic in their distribution among sponge specimens (Fig. 1B). These OTUs were most closely related to spirochetes detected in *Spongia officinalis* (OY759747.1) and *Astrosclera willeyana* (HE985144.1) sponges, respectively (Table S2). Inspection of phylogeny of these ten OTUs (Fig. 1C) revealed that six of the ten spirochete OTUs formed a clade with spirochete clones previously cloned from *T. favus* sponges (22). Of the remaining four, OTU105 and OTU128 (which were more abundant in non-Latrunculid sponge specimens) were part of distant clades of other sponge associated spirochetes, while OTU581 and OTU399 belonged to a clade stemming from a variety of environments (Fig. 1C). Notably, a clone (Sp02sw36) isolated from the seawater extruded from *Tsitsikamma favus* sponges in 2012 (22), was a close relative of the spirochetes associated with crown-of-thorns starfish (37), and the dominant spirochete found in *C. clathrus* sponges (33).

**Figure 1.**
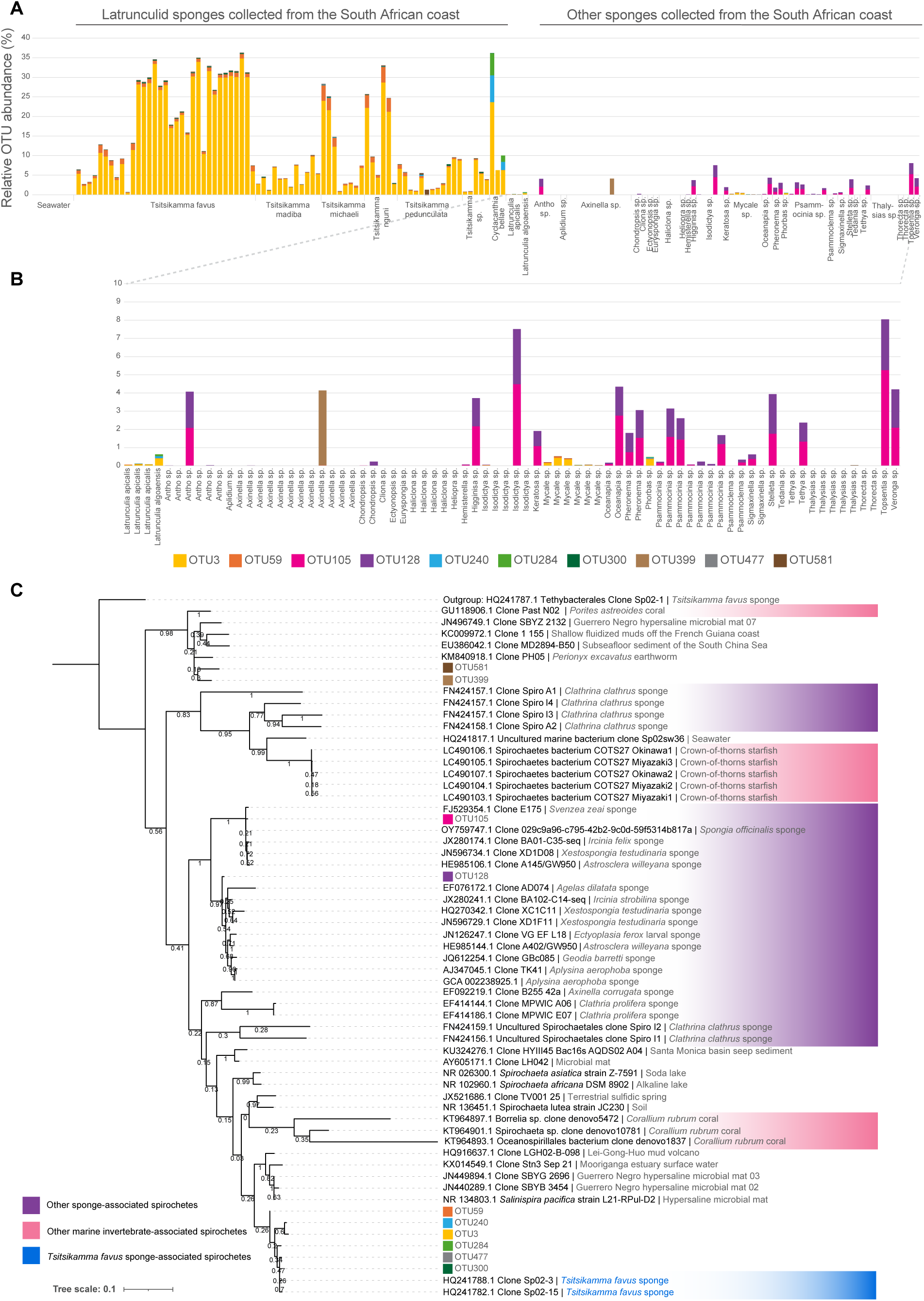
Spirochete population distribution in sponges collected from the South African coast and the Antarctic Southern Ocean. A) The relative abundance of OTUs clustered at a distance of 0.03 and classified as spirochetes, B) a magnified view of the spirochete OTUs present in non-latrunculid sponges collected from the south eastern coast of South Africa, three *L. apicalis* sponges collected from the Southern Ocean, and one sympatric *L. algoaensis* sponge. C) Maximum-likelihood phylogeny (with 1000 bootstraps) of the top ten most abundant spirochete OTUs recovered from the sponges included in this study.

### Characterization of *Tsitsikamma* sponge-associated spirochete MAGs

Eight sponges including five *T. favus* specimens (TIC2015-050A, TIC2015-050C, TIC2018-003B, TIC2018-003D, TIC2018-003M) and one each of *T. michaeli* (TIC2019-013N), *T. madiba* (TIC2022-009), and *T. pedunculata* (TIC2022-059) were selected for metagenomic analysis (Table S1). Following assembly, binning and taxonomic classification, eight spirochete MAGs were identified, one from each of the eight *Tsitsikamma* sponge metagenomes: MAGs 050A_2, 050C_7, 003B_7, 003D_7, 003M_1, 059_1, 013N_1, and 009_1 (Table 1, Table S3). The 16S rRNA and 23S rRNA gene sequences from each MAG (if recovered) were aligned against the NR nucleotide database via online BLASTn (46).

**Table 1.** Characteristics of putative representative genomes of *Tsitsikamma* sponge-associated spirochete symbiont MAGs.

| MAG | Size (Mbp) | Quality | 16S rRNA (% ID) | 23S rRNA (% ID) | Host | Sponge |
| --- | --- | --- | --- | --- | --- | --- |
| 003B_7 | 1.97 | Medium | N/A | *Salinispira pacifica* L21-RPul-D2 (89.54%) | *T. favus* | TIC2018-003B |
| 050A_2 | 2.73 | Low | Uncultured marine clone Sp02-3 (99.52%) | *Salinispira pacifica* L21-RPul-D2 (89.54%) | *T. favus* | TIC2016_050A |
| 003D_7 | 2.48 | High | Uncultured marine clone Sp02-3 (99.52%) | *Salinispira pacifica* L21-RPul-D2 (89.58%) | *T. favus* | TIC2018-003D |
| 003M_1 | 2.74 | High | N/A | *Salinispira pacifica* L21-RPul-D2 (89.58%) | *T. favus* | TIC2018-003M |
| 050C_7 | 1.72 | Medium | Uncultured marine clone Sp02-3 (99.52%) | N/A | *T. favus* | TIC2016-050C |
| 009_1 | 1.47 | High | N/A | *Salinispira pacifica* L21-RPul-D2 (91.25%) | *T. madiba* | TIC2022-009 |
| 013N_1 | 2.33 | High | N/A | *Salinispira pacifica* L21-RPul-D2 (89.48%) | *T. michaeli* | TIC2019-013N |
| 059_1 | 2.04 | Medium | N/A | N/A | *T. pedunculata* | TIC2022-059 |

The 16S rRNA gene sequences recovered from three MAGs all showed the greatest sequence identity with “Uncultured marine clone Sp02-3, ‘representing the conserved spirochete symbiont previously identified in *T. favus* sponges (22, 29). All recovered 23S rRNA sequences shared the greatest sequence similarity with *S. pacifica* L21-RPul-D2. This *S. pacifica* strain, isolated from a hypersaline microbial mat (47), was previously shown to be the closest known relative of the conserved spirochete Sp02-3 clone (22, 29). Finally, all eight *Tsitsikamma*-associated spirochete MAGs were taxonomically classified, via GTDB-Tk (48), within the *Salinispira* genus (Table S3). Therefore, we were confident these MAGs represented the conserved spirochete symbiont (Sp02-3) previously reported in South African latrunculid sponges.

### Phylogeny of *Tsitsikamma* sponge-associated spirochete MAGs

The 16S rRNA gene sequences recovered from three of the *Tsitsikamma*-associated spirochete MAGs were aligned against their closest matches in the NR database, and spirochetes from other marine invertebrates (37, 39), including the dominant spirochete present in the distantly related *Clathrina clathrus* sponges (33). Inferred maximum-likelihood phylogeny from the 16S rRNA gene alignment showed that the *Tsitsikamma*-associated spirochete MAGs were distinct from all other invertebrate-associated spirochetes (Fig. S1). The *Tsitsikamma*-associated spirochete MAGs formed a distinct clade but were most closely related to spirochetes detected in non-host-associated environments including hypersaline microbial mats, seawater, estuary water, and volcanic mud.

Since phylogeny inferred by a single marker gene can be limited, several orthogonal approaches were used to assess the phylogeny of the *Tsitsikamma* sponge-associated spirochete symbionts using whole genome data. Initially, we employed autoMLST (49) in *de novo* mode, with both concatenated alignment (Fig. 2A) and coalescent tree (Fig. 2B) approaches, using ten MAGs/genomes acquired from other sponge hosts, *Rhopaloides odorabile, Ircinia ramosa,* and *Aplysina aerophoba* (50–52), as references. The resultant phylogenies from these two approaches had largely congruent topologies, with the *Tsitsikamma* sponge-associated Sp02-3 symbionts and other sponge-associated spirochetes forming two related, but distinct clades (Fig. 2). The closest relative of the *Tsitsikamma*-associated spirochetes was *Salinispira pacifica*, in agreement with the 23S rRNA gene phylogeny. The *Tsitsikamma*-associated spirochetes appeared phylogenetically clustered following their respective hosts, rather than geographically clustered. This contrasted with other sponge-associated spirochetes that did not seem to follow any discernible pattern of possible co-phylogeny or phylosymbiosis (Fig. 2).

**Figure 2.**
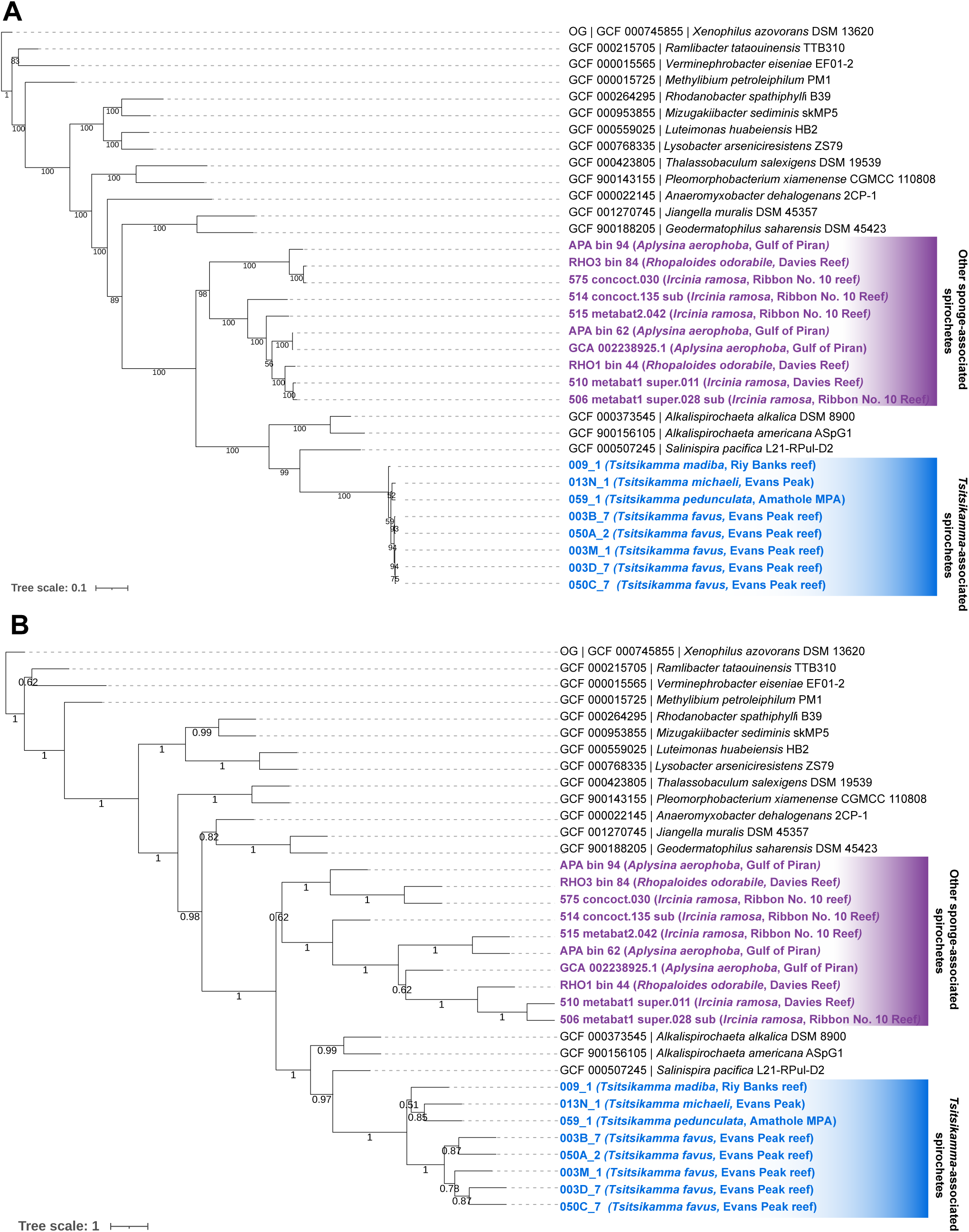
Phylogeny of sponge-associated spirochetes inferred with autoMLST in de novo mode using A) concatenated alignment and B) coalescent tree approaches. Tsitsikamma-associated spirochetes are highlighted in blue with their respective hosts. Other sponge-associated spirochetes are highlighted in purple with their associated hosts. All other reference spirochete genomes are listed in the format of ‘Accession number | Scientific name’.

As an orthogonal phylogenetic approach, we generated a phylogenetic tree using Phylophlan3 (53) and RaxML (54) (Fig. S2). Along with the eight *Tsitsikamma*-associated spirochete genomes and the ten genomes of spirochetes associated with other sponges, we included all Spirochaetaceae genomes from the NCBI database (N=300) and all host-associated spirochete MAGs from the JGI database (N=44). Again, the *Tsitsikamma*-associated spirochetes formed a clade distinct from all other sponge-associated spirochete genomes. Additionally, in this analysis, we found that a MAG present in seawater (GCA 913043885.1) clustered with the other sponge-associated spirochetes. The origin of this particular genome, whether from a free-living spirochete or a sponge symbiont, remains uncertain due to potential annotation errors in the database. However, we have opted to follow the supplied annotation and presume that this MAG is likely representative of the closest free-living relative within the clade. Our phylogenetic analysis incorporated all publicly available genomes and MAGs of the Spirochaetaceae phylum, and therefore this presumption is limited by the existing dataset. We calculated pairwise average nucleotide identity (ANI) scores for all 363 spirochete genomes (Table S4). The *Tsitsikamma*-associated spirochetes shared between 93.9% to 98.2% ANI with each other (Table S5), and less than 75% ANI with any other spirochete, including their closest relative *S. pacifica*.

### Estimated evolutionary divergence patterns of sponge-associated spirochetes

The divergence pattern of all sponge-associated spirochetes and their closest known free-living relatives was estimated using their rate of synonymous substitutions (dS) in orthologous genes present in all genomes. Visualization of the pairwise dS revealed that the *Tsitsikamma*-associated spirochetes are evolutionarily divergent from even their closest relative, *S. pacifica* (Fig. 3). It appears that the other sponge-associated spirochetes may have begun diverging before the *Tsitsikamma*-associated spirochetes diverged from their free-living relative. The divergence pattern of the *Tsitsikamma*-associated spirochetes is congruent with the phylogeny of their sponge host and incongruent with geographic location, suggestive of phylosymbiosis. Finally, it appears that these spirochetes have only recently begun diverging from one another as they adapt to their sponge host and that their association with latrunculid sponges is more recent than that of the co-dominant Tethybacterales symbionts (17).

**Figure 3.**
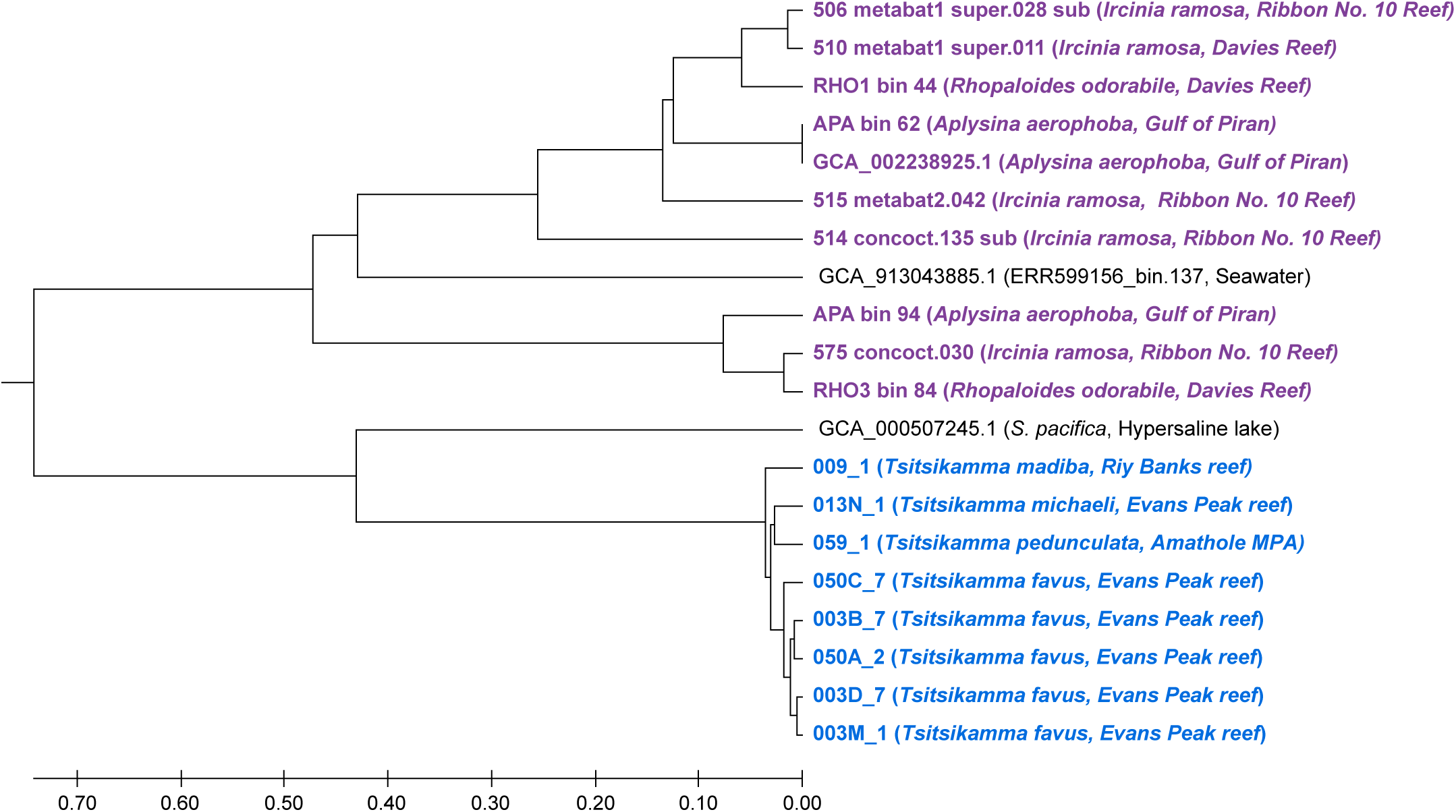
UPMGA representation of pairwise synonymous substitution rates (dS) of sponge-associated spirochete genomes, based on the alignment of 11 orthologous genes. PAL2NAL (88) and CodeML (89) from the PAML package were used to calculate pairwise dS values and the resultant matrix was visualized in MEGA11. The *Tsitsikamma*-associated spirochetes are colored in blue and other sponge-associated spirochetes are colored in purple.

### Comparative analysis of functional potential in spirochete genomes

The functional potential for all 363 spirochete genomes was predicted by assigning KEGG Orthologs (KO) annotations using KofamScan (55). KO counts per genome were mapped back to associated pathways detailed in the KEGG database (56) (Table S6). Dimension reduction of these counts per genome revealed distinct clusters suggestive of adaptation to the various environments from which these spirochetes were acquired (Fig. 4). The functional potential of the *Tsitsikamma*-associated spirochetes was distinct from spirochetes associated with other sponges and interestingly, was clustered more closely with the functional potential of spirochetes associated with oligochaete worms and spirochetes from hypersaline lake environments (Fig. 4).

**Figure 4.**
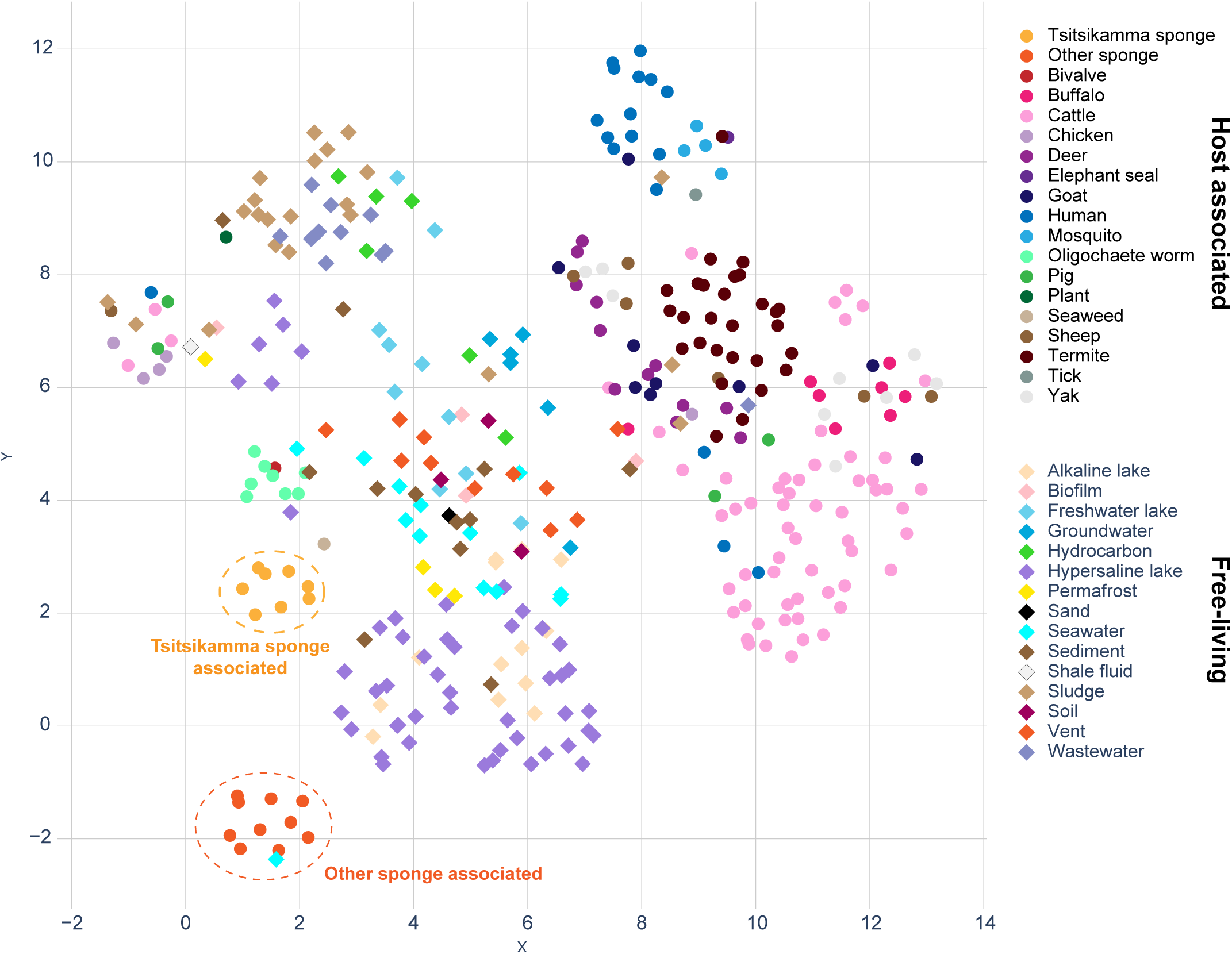
UMAP dimension reduction 2-dimensional representation of KEGG-annotated gene counts in all spirochete genomes. The isolation source of each genome is indicated by color, and shaped according to whether the isolation source is a living host (circles) or an abiotic environment (diamonds).

An Analysis of Similarity (ANOSIM) of the same data (Table S7) showed that the functional gene repertoire of the *Tsitsikamma*-associated spirochetes and other sponge-associated spirochetes were significantly different (*p* < 0.05) from one another and from all other environments. However, when considering the associated R-values, the *Tsitsikamma*-associated spirochetes may exhibit some overlap in functional potential of spirochetes in hypersaline lakes (R = 0.26), sediment (R = 0.31), freshwater lakes (R = 0.38), termites (R = 0.47), and seawater (R=0.49). This suggests that the functional repertoire of *Tsitsikamma*-associated spirochetes may be more akin to free-living species than host-associated.

### The biosynthetic potential of Sp02-3 spirochetes

A total of 581 biosynthetic gene clusters (BGCs) were detected in all spirochete genomes (N=363) (Table S8) and clustered into gene cluster families (GCFs) at a maximum distance of 0.3 with BiG-SCAPE (57) (Fig. 5A). Six of the eight *Tsitsikamma*-associated spirochetes had only a single predicted BGC. The remaining two MAGs, 003B_7 and 050A_2, which were of medium and low quality respectively, had no detected BGCs, likely due to incomplete coverage of the genomes. All six BGCs were predicted to encode a terpene product and were clustered into a single GCF (GCF1). Three other GCFs (GCFs 2, 3, and 4), consisting of terpene BGCs from other sponge-associated spirochetes, were identified but did not appear to have any homology with the terpene BGC in the *Tsitsikamma*-associated Sp02-3 spirochetes (Fig. 5B). Additional BiG-SCAPE analyses were performed with less stringent cutoffs of 0.5 and 0.8, and no BGCs from other spirochete genomes or the MiBIG database were incorporated into a GCF with the terpene BGCs detected in the *Tsitsikamma*-associated spirochetes (Table S8), indicating that this BGC is likely novel. Nonetheless, the closest characterized relative of the *Tsitsikamma*-associated spirochetes, *S. pacifica*, produces an orange carotenoid-like pigment (terpenoid), which we assume is produced via the only terpene BGC present in the *S. pacifica* genome. Despite the low sequence and organizational similarities, the terpene, if produced in the latrunculid-associated spirochetes, may protect them or their host against oxidative stress, as hypothesized for the *S. pacifica* bacterium (47)

**Figure 5.**
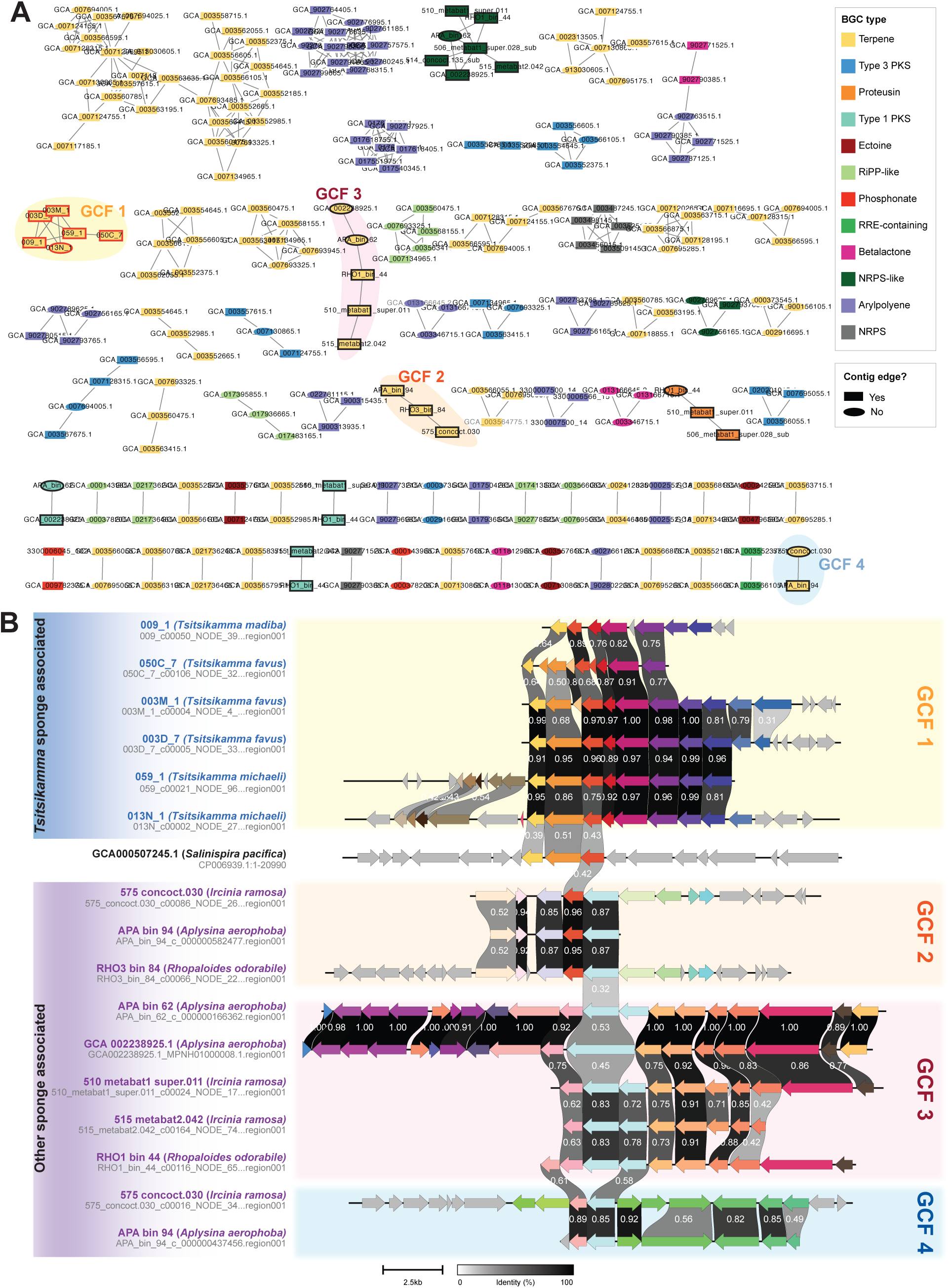
Assessment of biosynthetic potential in spirochetes. A) Network visualization of biosynthetic gene clusters from all spirochete genomes used in this study clustered into gene cluster families at a maximum distance of 0.3. BGCs from Tsitsikamma-associated spirochetes are highlighted with a red outline. BGCs from all other sponge-associated spirochetes are highlighted with a black outline. Gene cluster families (GCFs) of interest are highlighted. B) Pairwise comparison of amino-acid sequence identity of terpene biosynthetic gene clusters from sponge-associated spirochetes. The pairwise similarity between genes is indicated between genes, and genes are colored according to their predicted function. The GCFs to which the BGCs belong have been indicated.

In our previous studies, we have reported the existence of two chemotypes that exist in the *T. favus* and *T. michaeli* sponge populations in Algoa Bay (21, 28). Chemotype I represents the majority of sponges, as the sponges appear visually healthy with turgid structure and their spicules are in the canonical form. Further, this Chemotype is defined by the presence of a variety of discorhabdins and tsitsikammamines (28). Conversely, the morphology of the Chemotype II sponges is considered abnormal where the tissues appear bruised, are soft to the touch (akin to rotten fruit), and many spicules are malformed (28, 58). This chemotype is further characterized by the presence of structurally simpler makaluvamines and brominated discorhabdins (21, 28).

Previous surveys of the microbial communities associated with ten *T. favus* sponges and found no correlation between any bacterial population and the chemotypes (28). We have repeated the analysis with a larger cohort of *T. favus* and *T. michaeli* sponge specimens (N = 26). Using the same 16S rRNA gene amplicon datasets as presented in Figure 1, but instead including only data from the latrunculid sponges with associated chemical data, the analysis was repeated and OTUs were clustered at a maximum distance of 0.01 (Table S9) to disentangle the two spirochete strains previously identified in latrunculid sponges, Sp02-3 and Sp02-15 (22, 29). Using an Indicator Species Analysis (Table S10) we found that a decrease in Sp02-3 representative OTU abundance (OTU3) and an increase in Sp02-15 representative OTU abundance (OTU6) correlated with Chemotype II sponges, relative to Chemotype I specimens (Fig. S3 A – B, Table S10).

We conducted a correlation analysis of the top 50 most abundant OTUs with relative pyrroloiminoquinone abundance per sponge sample (Fig. S4, Table S11). The Sp02-3 spirochetes (OTU3) were positively correlated with the increased abundance of Chemotype I pyrroloiminoquinones and negatively correlated with the presence of Chemotype II pyrroloiminoquinones. The converse was true of the Sp02-15 spirochetes (OTU6) (Fig. S4, Table S11). As there was no evidence of BGCs for the production of pyrroloiminoquinones in the spirochete MAGs, this result suggests that the switch from Chemotype I to Chemotype II (the cause of which has yet to be identified) appears to negatively impact the Sp02-3 spirochete and allows the Sp02-15 spirochete to thrive in place.

Since the decrease in Sp02-3 similarly correlated with the incidence of deformed spicules, we considered whether it may play a role in spicule formation. The most closely related invertebrate-associated spirochete (Fig. 1 and Fig. 2) is a highly dominant and conserved spirochete in *Corallium rubrum* corals (39, 59) This spirochete is predicted to contribute to the coral’s overall health of the coral (60) and to produce a pigmented carotenoid that influences the commercially prized color of this red coral, as the spirochete’s presence correlates with the intensity of the observed red pigmentation (61). This spirochete was primarily found in the coenenchyme of the coral (61), which houses the sclerites (spicules) that are thought to act as initiation sites for the formation of the axial skeleton (62). Finally, the formation of spicules in a primary coral polyp is associated with a change in color from white to light pink (63). It is thus possible that the *C. rubrum*-associated spirochete may be involved in spicule formation as shown with the calcibacteria in *Hemimycale* sponges (pale orange to deep red in color) (64, 65), and hypothesized for the spirochetes in *Platygyra dadalea*, *Paragoniastrea australensis*, and Porites lutea sponges (44). While a speculative connection, as no MAG or genome is available for these spirochetes, this observation has prompted us to begin metatranscriptomic studies in conjunction with CARD-FISH experiments to determine the localization and potential structural role of spirochetes in latrunculid sponges from the South African coastline.

### Conclusion

This study shows that the conserved Sp02-3 spirochete of latrunculid sponges is likely to be a relatively new symbiont that has begun co-evolving with its respective sponge hosts. The Sp02-3 symbiont is distinct from all other invertebrate-associated spirochetes, including non-dominant spirochetes associated with other marine sponges. Assessment of their functional potential suggests that the Sp02-3 spirochetes are functionally unique relative to other sponge-associated spirochetes. We found no evidence that they are directly involved in the production of the pyrroloiminoquinones characteristic of their host sponges. The close phylogenetic relatedness of the latrunculid-associated spirochetes to a dominant, conserved coral-associated spirochete hints at a possibly structural role within the sponges. However, additional experiments will be necessary to test this hypothesis.

## METHODS AND MATERIALS

### Sponge Collection and taxonomic identification

Sponges were collected by SCUBA or Remotely Operated Vehicle (ROV) from multiple locations within the Tsitsikamma Marine Protected Area, Algoa Bay (Port Elizabeth), the Amathole Marine Protected Area (East London), and the Garden Route National Park. In addition, three *L. apicalis* specimens were collected by trawl net off Bouvet Island in the South Atlantic Ocean. Collection permits were acquired prior to collections from the Department of Environmental Affairs (DEA) and the Department of Environment, Forestry and Fisheries (DEFF) under permit numbers: 2015: RES2015/16 and RES2015/21; 2016: RES2016/11; 2017: RES2017/43; 2018: RES2018/44; 2019: RES2019/13; 2020: RES2020/31; 2021: RES2021/81; 2022: RES2022/70. Collection metadata are provided in Table S1. Sponge specimens were stored on ice during collection and moved to −20 °C on return to the lab. Subsamples of each sponge, collected for DNA extraction, were preserved in RNALater (Invitrogen) and stored at −20 °C. Sponge specimens were identified through inspection of gross morphology, spicule analysis, and molecular barcoding, as performed previously (21, 28, 29, 58).

### Bacterial community profiles in latrunculid sponges

The V4-V5 of the 16S rRNA gene was PCR amplified from 79 latrunculid sponges collected between 1994 and 2022 (See Table S1 for collection data). Amplicons were sequenced using the Illumina MiSeq platform and curated using mothur (v.1.48.0) (45). All raw amplicon read data can be accessed under accession number PRJNA508092. Briefly, sequences that were shorter than 250 nt in length, longer than 350 nt in length, had homopolymeric runs of 7 nt or more, had ambiguous bases, or had a sliding window quality average lower than 20, were removed from the datasets. Chimeric sequences were detected using VSEARCH (66) and removed from the dataset. Sequences were then classified via alignment against the SILVA database (v138.1) and any sequences classified as “Chloroplast”, “Mitochondria’’, “unknown”, “Archaea”, or “Eukaryota” were removed. Sequences were clustered into Operational Taxonomic Units (OTUs) at a distance of 0.03 and read counts thereof were converted to relative abundance (Table S2). Representative sequences of each OTU were aligned against the SILVA database (v138.1) in mothur and against the nt prokaryotic database using standalone blastn (67), using parameters-max_hsps 1-max_target_seqs 1 to return only the first match. Descriptions and isolation sources for each returned accession were retrieved using the esearch, efetch and xtract methods from the stand-alone entrez package (68). Spirochete OTUs were subset out and aligned with reference sequences from the NCBI nucleotide database using MUSCLE (v. 5.1) (69, 70) and phylogeny was inferred from the alignment using the Maximum-likelihood method with 1000 bootstrap replicates in MEGA11 (71). Finally, the same analysis was repeated but using only the raw amplicon read data from latrunculid sponges, and the OTUs were clustered at a distance of 0.01. in all other respects, the analyses were identical.

### Chemical Analysis and Chemotype Identification

Sponge extracts were prepared by extraction with methanol, drying *i. vac.* and resuspension in methanol at 1-10 mg/mL. LC-MS/MS data was acquired on a Bruker ESI-Q-TOF Compact (Bruker, Bremen) in positive ionization mode coupled to a Dionex Ultimate3000 Chromatograph (ThermoScientific, Sunnyvale, CA, USA) and using reversed-phase C18 columns and mobile phases consisting of water and acetonitrile with 0.1% formic acid each, using one of two methods (see Supplementary Methods for details). The data was converted to mzXML format and analyzed using MZmine3 (72) to assemble an aligned feature list (see Supplementary Methods for details). The feature list was filtered based on comparison of *m/z* values and MS/MS spectra to known or putative pyrroloiminoquinones. Peak area values were normalized to the overall pyrroloiminoquinone signal per sample and aggregated to the pyrroloiminoquinone class to summarize the latrunculid pyrroloiminoquinone profiles.

### Correlation of spirochete populations and sponge chemotypes

An Indicator species analysis was performed using the OTUs clustered at a distance of 0.01 for all *T. favus* and *T. micheali* sponges for which a chemotype had been assigned (16S_Chemotype_Indicator_Species_Analysis.R) to determine which OTUs, if any, were associated with the two chemotypes. The co-correlation analysis of the 50 most abundant OTUs (found as an average across all samples) was performed using the ‘cor’ function (73) native to R using dataframes of OTU and compound abundances as input. A 16S rRNA gene sequence phylogeny was built from the representative sequences of the top 50 OTUs, aligned with MUSCLE (v 5.1) (69, 70), using the neighbor-joining approach with 1000 bootstraps in MEGA11 (71). The final tree was visualized in iTol (74) where the correlation matrix and the average OTU abundance per sponge species was visualized alongside the tree as datasets.

### Metagenomic sequencing and analysis of individual *T. favus* specimens

The DNA extraction and metagenomic sequencing of four *Tsitsikamma favus* sponges that resulted in the recovery of four MAGs 050A_2, 050C_7, 003B_7, and 003D_7, classified as spirochetes, is described in Waterworth et al., 2021(17). In addition to these samples, four additional metagenomes of *Tsitsikamma* sponges (TIC2018-003M, TIC2019-013N, TIC2022-009, and TIC2022-059) were sequenced. These sponges were selected for sequencing based on the apparent abundance of spirochete OTUs found via 16S rRNA gene amplicon sequence.

Total genomic DNA (gDNA) was extracted using the Zymo Research Quick DNA Fecal/Soil Microbe Miniprep Kit (Catalog number: D6012) according to the manufacturer’s specifications and stored at −4 °C. Shotgun metagenomic IonTorrent libraries of 200 bp reads were prepared and sequenced using an Ion P1.1.17 chip. All metagenomes were assembled, binned, and processed as described in Waterworth et al., 2021 (17). Four additional spirochete genome MAGs (003M_1, 059_1, 013N_1, and 009_1) were extracted from the new datasets. MAGs were named after the *Tsitsikamma* sponge specimen from which they were extracted (e.g. 050A_2 is the MAG from sponge specimen TIC2016-050A). The numbers associated with each MAG are an arbitrary artifact of the binning process.

### Acquisition of reference genomes and MAGs

Four spirochete MAGs associated with *Aplysina aerophoba* and *Rhopaloeides odorabile sponges* from a study by Robbins and colleagues (75) were downloaded from https://data.ace.uq.edu.au/public/sponge_mags/, and five sponge-associated spirochete MAGs were acquired from the China National GeneBank DataBase (CNGBdb) from studies by O’Brien and colleagues (50, 51). One spirochete genome from an *Aplysina aerophoba* sponge was additionally downloaded from the NCBI database (GCA_002238925.1). Additionally, all other genomes classified within the Spirochaetaceae family were downloaded from the NCBI database (N=300) and all host-associated spirochete MAGs were downloaded from the JGI database (N=44). This resulted in a total of 354 reference genomes (Table S3).

### Characterization of MAGs and genomes

All scripts used for bioinformatic analyses, and their associated inputs, used in the following methods can be found at https://github.com/samche42/Spirochete. All MAGs and genomes used in this study were assessed using CheckM (v1.1.3) (76) and taxonomically classified using GTDB-Tk (v2.3.2) (48) against the Release 214.1 reference database. Basic metrics such as size, number of contigs, and N50 were calculated using bin_summary.py. The number of genes, pseudogenes, and coding density per genome were calculated using all_included_genome_characteristics.py. All metadata per genome or MAG can be found in Table S3.

### Phylogeny of spirochete genome MAGs extracted from individual *Tsitsikamma* sponges

Ribosomal sequences (23S rRNA, 16S rRNA, and 5S rRNA) were extracted from individual MAGs using barrnap (v 0.9) (77). The closest matches of recovered 16S sequences from sponge-associated MAGs were identified using BLASTn (v 2.7.1) (67). Resultant sequences were aligned using MUSCLE (v. 5.1) (69, 70) and phylogeny was inferred using the Maximum-likelihood method with 1000 bootstraps in MEGA11 (71). Phylogeny of the *Tsitsikamma*-associated spirochete MAGs was similarly inferred using whole genome data via autoMLST (49) and PhyloPhlan3 (53). Amino acid sequences and nucleotide sequences for all genes were found in all genomes using prokka (v 1.13) (78). The phylogeny of all 362 MAGs and genomes (8 *Tsitsikamma*-associated spirochete MAGs and 354 references) was inferred using Phylophlan3: Phylophlan3 was run with diversity set to medium, with default values in the supermatrix_aa configuration. The resultant gene protein alignment was used in RaxML (v 8.2.12) (79) to build a phylogenetic tree with 1000 bootstrap replicates using the PROTGAMMAAUTO model. The resultant tree was visualized in iTol (74). Genomes from Myxococcota (GCA_002691025.1) and Deltaproteobacteria (GCA_020632655.1) were chosen as outgroups. These genomes had been downloaded from the NCBI database as their metadata indicated that they were classified within the Spirochaetaceae family. However, the taxonomic classification of these genomes with GTDB-Tk revealed that these genomes had likely been misclassified. These genomes were considered serendipitous choices for outgroups for the Phylophlan3 analysis. AutoMLST was deployed in *de novo* mode using concatenated alignments and coalescent trees of marker genes in two separate analyses. ModelFinder and IQ-TREE Ultrafast Bootstrap analysis were enabled in both analyses. All latrunculid-associated and other sponge-associated spirochete MAGs were included in this analysis. MAGs and genomes from JGI and NCBI were not used in this analysis as the number of query genomes is limited to 20 so we opted to include only sponge-associated spirochetes in this analysis. Resultant trees were downloaded in Newick format and revisualized in iTol (74). Finally, the pairwise average nucleotide identity (ANI) was calculated for all genomes using fastANI (v1.33)(80). If a pairwise alignment fraction (AF) was lower than 70% (81), the associated ANI score was nullified as the accuracy of the ANI score could not be trusted.

### Estimated evolutionary divergence patterns of sponge-associated spirochetes

Using the Phylophlan3 (53) and autoMLST(49, 53) trees as guidance, orthologous genes from the eight *Tsitsikamma*-associated spirochetes, the ten other sponge-associated spirochetes, and their closest relatives were identified using OMA (v. 2.6.0) (82). A total of 11 orthologs common to all genomes were found using count_OGs.py and aligned using MUSCLE (v 5.1) (69, 70). The corresponding nucleotide sequence for each gene was retrieved using streamlined_seqretriever.py, all stop codons were removed using remove_stop_codons.py, and nucleotide sequences were aligned using MUSCLE (v 5.1) (69, 70). Ortholog gene sequences were grouped per genome using merge_fasta_for_dNdS.py. The nucleotide and amino acid sequences (per genome) were each concatenated union function from EMBOSS (83) and aligned using PAL2NAL (84). The alignment was used to estimate pairwise synonymous substitution rates (dS) and thereby infer the pattern of divergence between these genomes using codeml from the PAML package (85).

### Comparative analysis of functional potential in spirochete genomes

Genes were identified in all genomes/MAGs using Prokka (v 1.13) (78) and then annotated against the KEGG database using KOfamSCAN (55) with detail-tsv as the output format. Reliable annotations were extracted from these results based on the criteria that the annotation score is greater than the estimated threshold, and then reliable annotations per MAG/genome were counted and summarized using the kegg_parser.py script. This produced a table of KO counts per genome that was used as input for both Analysis of Similarity (ANOSIM.R) processing and dimension reduction, via UMAP (86), for 3-dimensional and 2-dimensional visualizations (dimension_reduction.py). A Jupyter notebook is provided in the GitHub repository for easy reproduction and an interactive 3D figure. To find statistically significant KEGG-annotated drivers of the different samples, we performed a re-purposed Indicator Species Analysis with the number of KEGG annotations per KO per genome in place of OTU abundance. This was performed using the multiplatt method from the “indicspecies” package in R (87) with 1000 permutations and specifying the point biserial correlation coefficient (“r.g”) as the association index as this both accounts for abundance data (rather than presence/absence data) and corrects for the different number of samples per host type.

### The biosynthetic potential of sponge-associated spirochetes

A total of 547 biosynthetic gene clusters (BGCs) were predicted from all spirochete genomes (N=363) using antiSMASH (v. 6.0.1) (88) with--cb-general--cb-knownclusters--cb-subclusters--asf--pfam2go--smcog-trees options enabled and genes found with prodigal. The resultant putative BGCs were clustered twice using BiG-SCAPE (v 1.1.5)(57) at maximum distances of 0.3, 0.5, and 0.8. Network files of non-singleton gene cluster families (GCFs) were visualized in Cytoscape (89). Highlighted gene clusters of interest were visualized with clinker (90). Metadata for BGCs was extracted from individual GenBank files using antismash_summary.py.

## DATA AVAILABILITY

All sequence data can be accessed under accession number PRJNA508092 in the NCBI SRA database. All scripts used for analysis and visualization can be accessed at https://github.com/samche42/Spirochete.

## ACKNOWLEDGEMENTS

We would like to acknowledge Gwynneth Matcher (South African Institute for Aquatic Biodiversity, Aquatic Genomics Research Platform), Carel van Heerden and Alvera Vorster (Stellenbosch University Central Analytical Facility) for next-generation sequencing technical support. We thank Ryan Palmer and Koos Smith (African Coelacanth Ecosystem Programme) for logistics and technical support during sponge collections. We thank the South African Environmental Observation Network, Elwandle Coastal Node, and the Shallow Marine and Coastal Research Infrastructure for the use of their research platforms and infrastructure for their assistance in SCUBA collections and logistical support. This research was supported by South African National Research Foundation grants to R.A.D., including the South Africa Research Chair Initiative (SARChI) grant (UID: 87583) and the SARChI-led Communities of Practice Programme (UID: 110612). S.C.W. was supported by an NRF Innovation and Rhodes University Henderson Ph.D. scholarships. G.M.S. and L.M. were supported by NRF Masters and PhD scholarships, respectively. S.P.-N. Was supported by an NRF PDP scholarship (UID: 101038). J.C.J.K. was supported by funding awarded to R.A.D. by the South African Medical Research Council as well as the UK Medical Research Council, with funds received from the UK Government’s Newton Fund (Grant No.: 96185). We declare no competing interests, financial or otherwise, in relation to the work described here. The opinions expressed and conclusions arrived at are those of the authors and are not necessarily to be attributed to any of the above-mentioned donors.

