## Supplementary material for "The unique and enigmatic spirochete symbiont of latrunculid sponges": Fig. S2

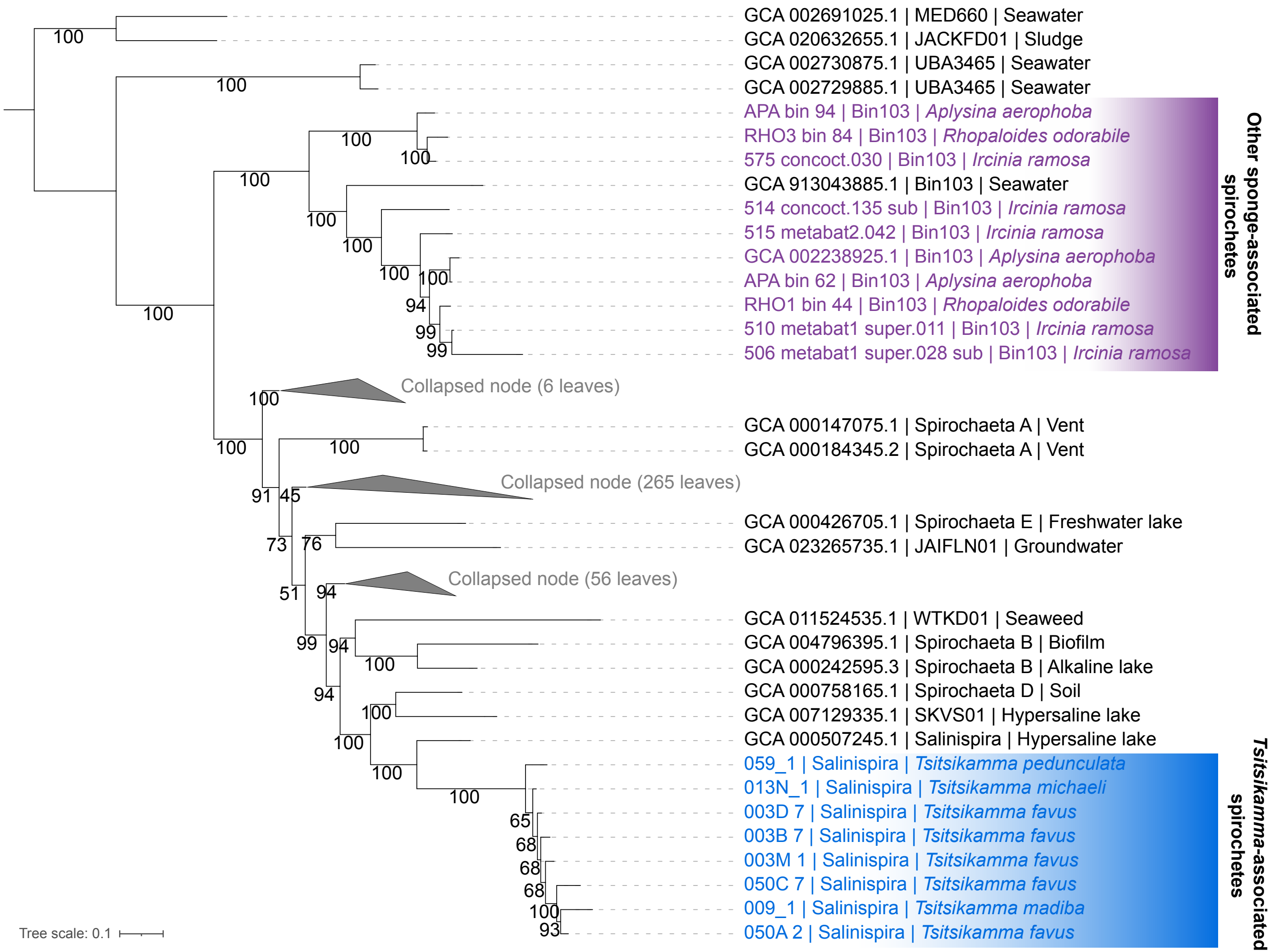


**Figure S2.** Phylogeny of all Spirochaetaceae genomes as inferred using an alignment generated by Phylophlan3 and RaxML with 1000 bootstrap replicates. Tsitsikamma-associated spirochetes are highlighted in blue, and other sponge-associated spirochetes are highlighted in purple. Bootstrap values and branch length are indicated.
