## Supplementary material for "The unique and enigmatic spirochete symbiont of latrunculid sponges": Fig. S3

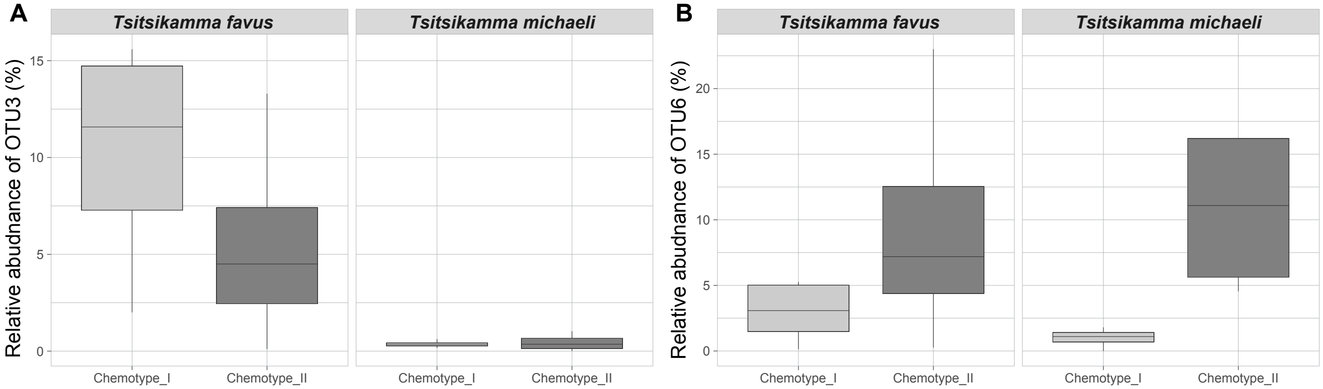


**Figure S3.** The distribution of relative abundance of A) OTU3 and B) OTU6 in Chemotype I and Chemotype II sponges from the T. favus and T. michaeli species
