## Supplementary material for "The unique and enigmatic spirochete symbiont of latrunculid sponges": Fig. S4

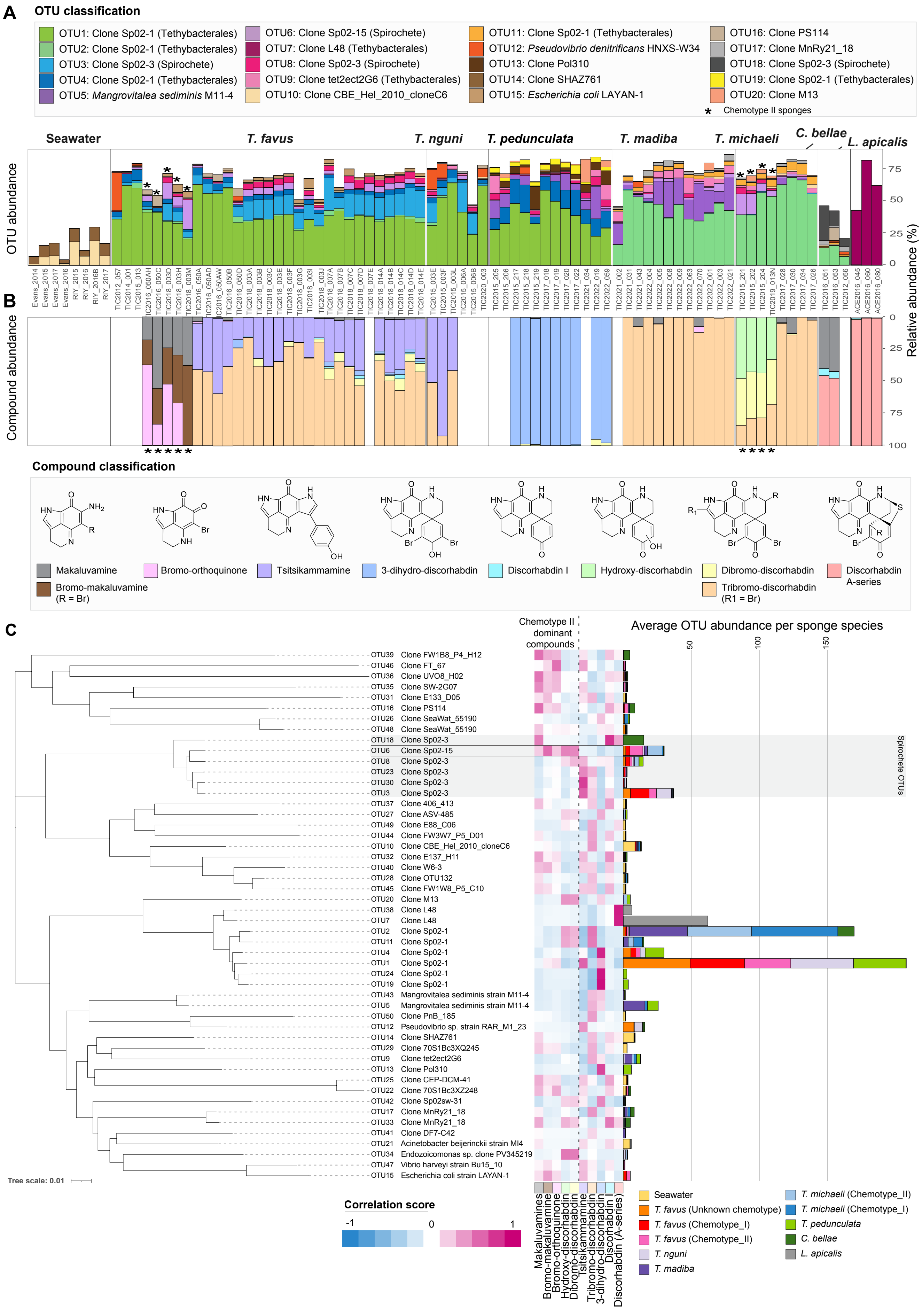


**Figure S4.** Distribution of microbial OTUs and pyrroloiminoquinone classes in latrunculid sponges and the correlation thereof. A) Bacterial community profiles associated with latrunculid sponges and surrounding seawater represented by OTUs clustered at a distance of 0.01. The relative abundance of the top 20 OTUs with the highest average abundance across all samples is shown. The closest relative determined by alignment against the NCBI nt database is indicated within the colored key legend. B) The relative abundance of different pyrroloiminoquinone classes per sponge sample as indicated by the colored key. Scaffold structures are provided for identified chemical groups. Blank regions indicated sponges that have not had their chemistry assessed. Sponges classified as Chemotype II have been indicated with an asterisk. Sponge samples from which metagenomic data was retrieved and analyzed are indicated with bold lettering. C) Co-correlation analysis of OTU and pyrroloiminoquinone compound abundance shows that OTU6 is exclusively correlated with all compounds that are characteristic of T. favus Chemotype II and T. michaeli Chemotype II sponges. The correlation score per OTU-compound combination is indicated by a color scale ranging from -1 to 1. A negative score is indicative of a presence-absence relationship and a positive score is indicative of a presence-presence relationship.
