## Supplementary material for "The unique and enigmatic spirochete symbiont of latrunculid sponges": Fig. S1

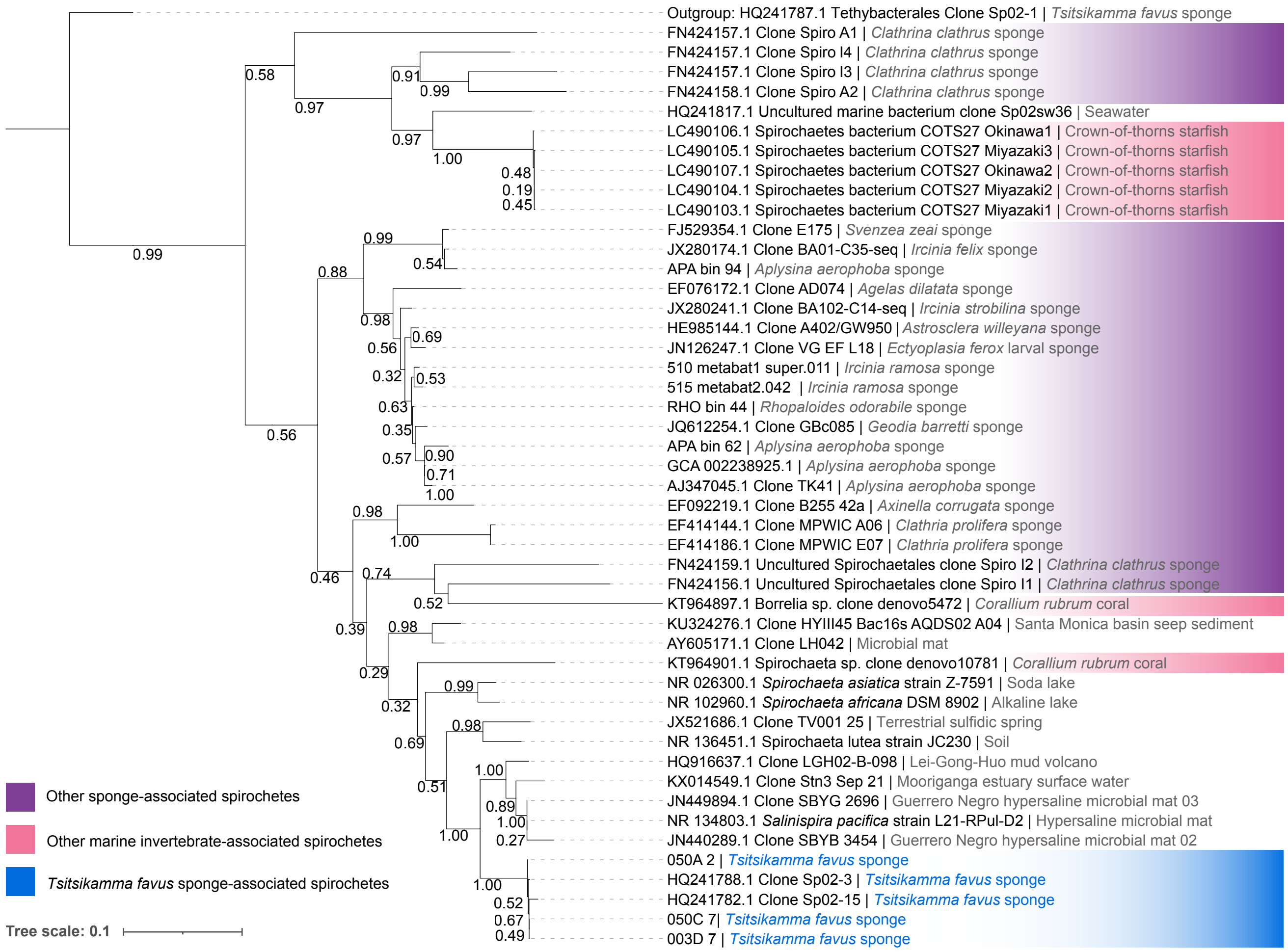


**Figure S1.** Maximum-likelihood phylogeny of spirochete 16S rRNA gene sequences inferred with 1000 bootstrap replicates. A 16S rRNA gene sequence from a sympatric Tsitsikamma-associated Tethybacterales clone was used as an outgroup. Bootstrap values are indicated on branch nodes. Sequences from latrunculid-associated spirochetes are indicated in blue, other sponge-associated spirochetes are indicated in purple, and spirochetes associated with other marine invertebrates are indicated in pink. For all sequences, accessions are provided where available, along with their description per the NCBI database and isolation source.
